# A Random Forest Classifier Uses Antibody Responses to *Plasmodium* Antigens to Reveal Candidate Biomarkers of the Intensity and Timing of Past Exposure to *Plasmodium falciparum*

**DOI:** 10.1101/2022.02.16.480705

**Authors:** Sophie Bérubé, Tamaki Kobayashi, Douglas E. Norris, Ingo Ruczinski, William J. Moss, Amy Wesolowski, Thomas A. Louis

## Abstract

Important goals of malaria surveillance efforts include accurately quantifying the burden of malaria over time, which can be useful to target and evaluate interventions. The majority of malaria surveillance methods capture active or recent infections which poses several challenges to achieving malaria surveillance goals. In high transmission settings, asymptomatic infections are common and therefore accurate measurement of malaria burden often demands active surveillance; in low transmission regions where infections are rare accurate surveillance requires sampling a large subset of the population; and in any context monitoring malaria burden over time necessitates serial sampling. Antibody responses to *Plasmodium falciparum* parasites persist after infection and therefore measuring antibodies has the potential to overcome several of the current difficulties associated with malaria surveillance. However, identifying which antibody responses are markers of the timing and intensity of past exposure to *P. falciparum* is challenging, particularly among adults who tend to be re-exposed multiple times over the course of their lifetime and therefore have similarly high antibody responses to many *P. falciparum* antigens. A previous analysis of 479 serum samples from individuals in three regions in southern Africa with different historical levels of *P. falciparum* malaria transmission (high, intermediate, and low) revealed regional differences in antibody responses to *P. falciparum* antigens among children under 5 years of age. Using a novel bioinformatic pipeline optimized for protein microarrays that minimizes between-sample technical variation, we used antibody responses to *P. falciparum* and *P. vivax* antigens as predictors in random forest models to classify adult samples into these three regions of differing historical malaria transmission with high accuracy. Many of the antigens that were most important for classification in these models do not overlap with previously published results and are therefore novel candidate markers for the timing and intensity of past exposure to *P. falciparum*. Measuring antibody responses to these antigens could potentially lead to improved malaria serosurveillance that captures the timing and intensity of past exposure.

## 2 Introduction

Accurately quantifying the burden of malaria is an important component of malaria control and elimination efforts (The World Health Organization, 2020) allowing more targeted interventions to be deployed and evaluated (Moonen *and others*, 2010). Currently, common malaria surveillance methods include measuring incidence or prevalence of malaria infections based on active or recent infections using diagnostic tests (e.g. microscopy, rapid diagnostic tests, or polymerase chain reaction). However, there are challenges associated with using these methods. In low transmission settings incident infections are rare, and in high transmission areas asymptomatic infections are common; both factors make detecting active or recent infections resource intensive (Mueller *and others*, 2011; Cibulskis *and others*, 2011; Satoguina *and others*, 2009). Additionally, surveillance that detects active or recent infections provides estimates of malaria burden at single point in time. While this information is useful, collecting these data across multiple time points is essential to answer important scientific questions such as how malaria transmission changes over time, following interventions, and seasonally. With existing methods, surveillance over time requires sampling systems like household surveys or reactive case detection to be sustained for long periods; a resource intensive demand (Camargo *and others*, 1999; Trape *and others*, 2014). Antibody responses to *Plasmodium falciparum* infection have the potential to overcome several of these challenges because antibodies persist for extended periods after infection and can provide insight into an individual’s or a population’s infection history without needing to capture active infections. Serological data can also provide longitudinal information about malaria burden with a cross-sectional sample, which requires substantially fewer resources to obtain (Corran *and others*, 2007; Drakeley and Cook, 2009).

Interpreting antibody responses to *P. falciparum* infections is complicated by several factors including: multiple developmental phases of the parasite in humans, each with different expressed genes that produce various antigens; the presence of multiple genetically distinct clones in a single infection, possibly eliciting different antibody responses; and reinfection events which are common over short time scales in high transmission settings (Hafalla *and others*, 2011; Gonzales *and others*, 2020; Felger *and others*, 2012). Despite these challenges, studies have illuminated many aspects of the human antibody response to *P. falciparum* infections, including correlates of protective immunity, possible vaccine targets (Cockburn and Zavala, 2007; Trieu *and others*, 2011), and as markers of past infection (Helb *and others*, 2015; Crompton *and others*, 2010). However, such studies have largely been successful in children under the age of 5 years, and expanding these studies to broader age groups is complicated by the fact that older individuals in malaria endemic settings tend to be re-exposed and re-infected multiple times throughout their life. These infection events give adults across regions of differing transmission similarly high reactivity to *P. falciparum* antigens Helb *and others* (2015); Kobayashi *and others* (2019); van den Hoogen *and others* (2020), and this makes distinguishing antibody responses based on differences in past exposure difficult. The ability to distinguish the timing and intensity of adults’ past exposure to malaria is essential for long-term surveillance goals, since it allows analysts to make inferences about past malaria transmission levels over a longer time frame. In many settings, only adults have had sufficient repeated exposure to acquire immunity to clinical malaria, and thus make up the majority of the asymptomatic reservoir which is thought to be disproportionately responsible for onward transmission (Bousema *and others*, 2014; Sumner *and others*, 2021). Therefore, serology based surveillance methods that can be applied to adults have the potential to inform more effective interventions since this information may better characterize the asymptomatic reservoir.

In order to expand the search for markers of the intensity and timing of past exposure to *P. falciparum* in adults, we used novel bioinformatic tools to analyze protein microarray data measuring antibody responses to 500 *P. falciparum* and 500 *P. vivax* antigens in 479 serum samples from individuals aged 0 − 86 years in three regions in Zambia and Zimbabwe (Kobayashi *and others*, 2019). These three regions span low, intermediate and high historical malaria transmission levels (Moss *and others*, 2015; Mharakurwa *and others*, 2012). If antibody responses to certain *Plasmodium* antigens predict an individual’s region of residence with high accuracy across multiple age groups, these could be indicative of differences in the frequency of past exposure over decades, which is highly dependent on the intensity of transmission over this time frame. Additionally, the probability of recent exposure is directly correlated to the level of transmission and, therefore, antibody responses that are relatively constant across regions of differing transmission levels are unlikely to be markers of recent exposure. Consequently, antibody responses that differentiate individuals residing in regions of differing historical malaria transmission could be markers of the intensity and timing of past exposure to *P. falciparum* over a long time frame. Measuring these antibodies could overcome the current challenges associated with malaria surveillance.

A previous analysis of these data published by Kobayashi *and others* (2019) revealed a correlation between region of origin and antibody responses to a subset of the 30 *P. falciparum* most seroreactive antigens among children under 5 years of age, but little correlation was observed among older individuals. Here, we conduct an analysis of these data with a novel bioinformatic pipeline (Bérubé *and others*, 2021, 2022) that ranks antibody responses to each antigen within each sample relative to that individual’s response to other *Plasmodium* antigens thereby minimizing the effects of between-sample technical variation, and thus making it more feasible to distinguish adult antibody responses, which tend to be more uniformly high across all transmission settings. These ranks are then used as predictors for a random forest model that classifies samples into their region of origin. Using subsets of 20 to 260 antigens, we classify all samples, including those from adults, into their region of origin with high accuracy. The random forest approach generally out-performs parametric multivariate modelling (e.g., logistic regression), particularly when many antigens were used to predict the location of origin of a sample. This superiority suggests models that leverage between-antigen interactions play an important role in using antibody responses to differentiate exposure to varying levels of malaria transmission. A closer analysis of the antigens most important for classification reveals some overlap with previously identified markers of recent past exposure, however, several identified antigens appear to be novel markers of the intensity and timing of past exposure and so potentially useful tools for malaria surveillance.

## 3 Materials and Methods

### 3.1 Study populations and sites

A total of 479 samples from Choma District, Zambia, Nchelenge District, Zambia, and Mutasa District, Zimbabwe were probed with protein microarrays containing 500 *P. falciparum* and 500 *P. vivax* antigens. Of these, 414 samples were collected in serial, cross-sectional, community-based surveys in 2015 across all three sites, including at health centers and 65 samples from 13 individuals residing in Choma District, Zambia were collected from 2013-2015 in a longitudinal study. Across the three locations, the proportion of samples positive for *P. falciparum* by rapid diagnostic test (RDT) at the time of sampling, was between 3 and 35% and the proportion of individuals who were febrile (temperature ≥ 38° C) at the time of sampling ranged from 5 to 21%. Additionally, ages of individuals at the time of sampling are comparable across all three sites and span a wide range (0 − 86 years) (Table 1) (Kobayashi *and others*, 2019).

**Table 1:** Study population and site characteristics.

| Site | Choma District, Zambia | Nchelenge District, Zambia | Mutasa District, Zimbabwe |
| --- | --- | --- | --- |
| Number of Samples | N=166 | N=122 | N=190 |
| Mean Age (Range) in years | 32.8 (1-83) | 23 (1-65) | 15 (0-86) |
| % Female | 43 | 61 | 56 |
| Average Weekly RDT Positive Cases at Health Centers |  |  |  |
| 2012 | 1.0 | 100.0 | 24.4 |
| 2013 | 0.7 | 98.3 | 28.8 |
| 2014 | 1.3 | 146.0 | 24.2 |
| 2015 | 0.9 | 115.0 | 6.9 |
| 2016 | 3.1 | 74.0 | 8.4 |
| % RDT Positive Samples | 21 | 35 | 3 |
| % Febrile at time of Sampling | 21 | 13 | 5 |

Although the characteristics of individuals across the three sites are similar, the levels of historical malaria transmission differed greatly. Table 1 shows the average weekly number of RDT positive cases at health centers in each of the three regions from 2012 − 2016. These numbers reveal a consistently low prevalence of malaria in Choma District, Zambia, a consistently high prevalence of malaria in Nchelenge District, Zambia and an intermediate level of transmission from 2012 − 2014 followed by a sharp decline in 2015 in Mutasa District, Zimbabawe. These figures are compiled from data published in Moss *and others* (2015); Mharakurwa *and others* (2012), and are representative of malaria transmission levels in these regions from 2012 − 2016. All cases of malaria in these regions are caused by *P. falciparum* and *P. vivax* is not known to have been present in any of these three areas from 2012 − 2016, or currently.

### 3.2 Pre-Processing Measurements from Protein Microarrays

Output from the pre-processing pipeline described in Bérubé *and others* (2021) is fed into a Bayesian model that produces full posterior distributions of the true, underlying protein signal. Briefly, for pre-processing, the ratio of observed foreground signal to background signal at each probe or antigen (*Y^′^*) is log transformed. Then, using a robust linear model described by Sboner *and others* (2009), array and subarray effects are estimated and subtracted from log(*Y^′^*) to produce 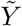. Finally, these values are standardized using array specific sample means and sample standard deviations of control probes to produce *Y*, the input for the Bayesian model (Equation 1).

We denote measurements from microarrays *Y*_*i,p*_ with the following subscripts:

- array number *i* ∈ {1, …, *I*}, since each array corresponds to a single sample, *I* = 479 in this study.
- Antigens *p* ∈ {1, …, *P*}, in this study *P* = 1000, each *p* is a *Plasmodium* antigen on the array.

As described in Bérubé *and others* (2022), the Bayesian model (Equation 1) uses the transformed data, *Y*, to produce the full posterior distribution of the true underlying signal, denoted by *S*. We assume that true underlying fluorescent intensity resulting from specific protein binding between the proteins in the sample and the probes on the array, *S*, adds to a term *e*, which represents the technical variation or biological variation that is not the result of differences in exposure to *P. falciparum* remaining in the measurements after the pre-processing pipeline (Bérubé *and others*, 2021), to produce the normalized observation *Y*. We leverage control probes on the array to estimate the error distribution *e*, and ultimately produce an estimate of true underlying signal given observed and pre-processed signal, or *S*|*Y*, for each antigen on the array; this estimate is a full posterior distribution. The full hierarchical Bayesian model describing this relationship is described in Bérubé *and others* (2022).

A simplified version of the Bayesian model is as follows:

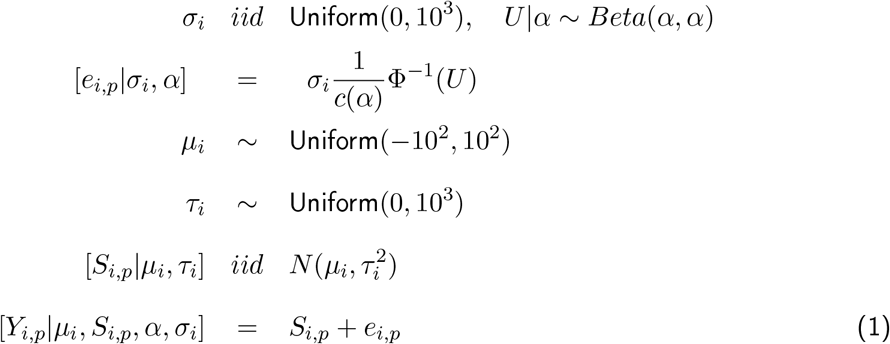

### 3.3 Ranking Pre-Processed Measurements

We compute the ranks of the true *S* at each antigen *i* on the array via:

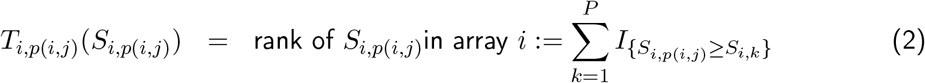

so that the smallest *S* will have rank 1 and the largest will have rank *P*. The estimator that minimizes squared error loss is the posterior mean:

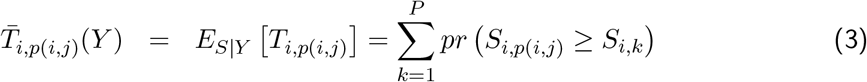

We use burned in Markov Chain Monte Carlo (MCMC) draws (Gelman *and others*, 2009) to compute this quantity by computing the rank of each protein in each array for each MCMC draw, and then averaging those ranks across the MCMC draws.

### 3.4 Random Forest Models

#### 3.4.1 Antigen Selection

We first fit a random forest model using the R package randomForest (Liaw and Wiener, 2002) using as predictors the ranks 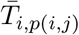 of all 1038 active probes (500 *P. falciparum* or *P. vivax* antigens expressed in *E. coli* and 38 purified *P. falciparum* antigens) on all 479 arrays spotted with samples from individuals residing in Zambia and Zimbabwe (described in Section 3.1). The random forest classified samples into their region of origin (Choma District, Zambia; Nchelenge District, Zambia; and Mutasa District, Zimbabwe). The mean decrease in Gini index is the metric used for determining variable importance. The top 20, 30, 65, 130, and 260 antigens are therefore those with the greatest mean decrease in Gini index as computed from this original model. After considering models fit to all 479 samples with the top 20, 30, 65, 130, and 260 antigens, we also consider models fit to subsets of samples using the top 20, 30, 65, 130, and 260 antigens as predictors. These subsets include children and adults (using a cutoff age of 5 years, and a cutoff of 15 years), and samples from individuals who are negative by rapid diagnostic test (RDT) at the time of sampling. For all random forest models we used a prior distribution with equal weights for each of the three classes (regions), the number of candidate variables selected randomly at each split is taken to be 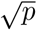, where *p* is the number of predictors in the model, and the number of trees to grow is set at 500.

#### 3.4.2 Assessing Model Performance

Models fit with either subsets of antigens as predictors or subsets of samples are evaluated using both out of bag (OOB) error (Liaw and Wiener, 2002; Breiman, 2001), and multiclass AUC with corresponding 95% confidence intervals are computed using the R package multiRoc (Wei and Wang, 2018). Specifically the global multiclass AUC is computed using the macro-average method of class-specific AUC as described by Hand and Till (2001).In addition to the OOB error and AUC computed by fitting random forest models to all 479 samples, we also evaluate the performance of models with a 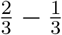 train-test split, and compute a test error, as well as an AUC with the reserved 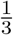 test data. All relevant R code to execute model fitting and checks can be found at https://github.com/sberube3/Antigen_classification_code.

## 4 Results

### 4.1 Responses to *Plasmodium* antigens are predictive of an individual’s region of residence

First, protein microarray data from all 479 serum samples were pre-processed and within-sample ranks (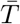, Section 3.3) of the relative reactivity to all antigens were computed based on fluorescent intensity. These values were then used as predictors in random decision forest models that classified all 479 serum samples into their geographical region of origin (Choma District, Zamiba, Nchelenge District, Zambia, or Mutasa District, Zimbabwe). In order to fine-tune our models we measured variable importance and defined top antigens to be those those with the highest mean decrease in Gini index from a random forest model with all 1038 antigens as predictors. We evaluated models with the top 20, 30, 65, 130, and 260 antigens as predictors and found that all of these models had global out of bag (OOB) error rates below 10% (Figure 1) and global AUC values above 0.95 (Table S1). Similar results were obtained by computing test error (Figure S1) and AUC (Table S2) from a 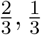 test-training split of the data, and using a random set of 20, 30, 65, 130, or 260 antigens as predictors produced consistently higher OOB error rates (Figure 1), reinforcing that the top 20 − 260 antigens accurately predict region of origin. Models that used fewer than the top 10 antigens as predictors had substantially higher cross-validated and OOB error rates (Table S6), suggesting that information contained in the ranks of the top 10 − 20 antigens contribute substantially to differentiating antibody responses across the three regions. By contrast, performance of the random forest models using the top 20, 30, 65, 130, and 260 antigens as predictors are highly comparable (Figure 1, Table S1, Table S6), suggesting that adding more than 20 top antigens does not substantially improve the accuracy of the models. These patterns are consistent for global OOB error rates, and across all three region-specific error rates. Since malaria transmission levels across these three regions spans a wide range of historic and current transmission intensities (Table 1), it follows that the exposure of people in this sample to *P. falciparum* parasites also spans high, intermediate and low levels, both historically and at the time of sampling. Therefore, the ability of these random forest models classify samples by region of origin with high accuracy suggests that antibody responses to a subset of these antigens could be markers of the intensity and timing of past *P. falciparum* exposure.

**Figure 1:**
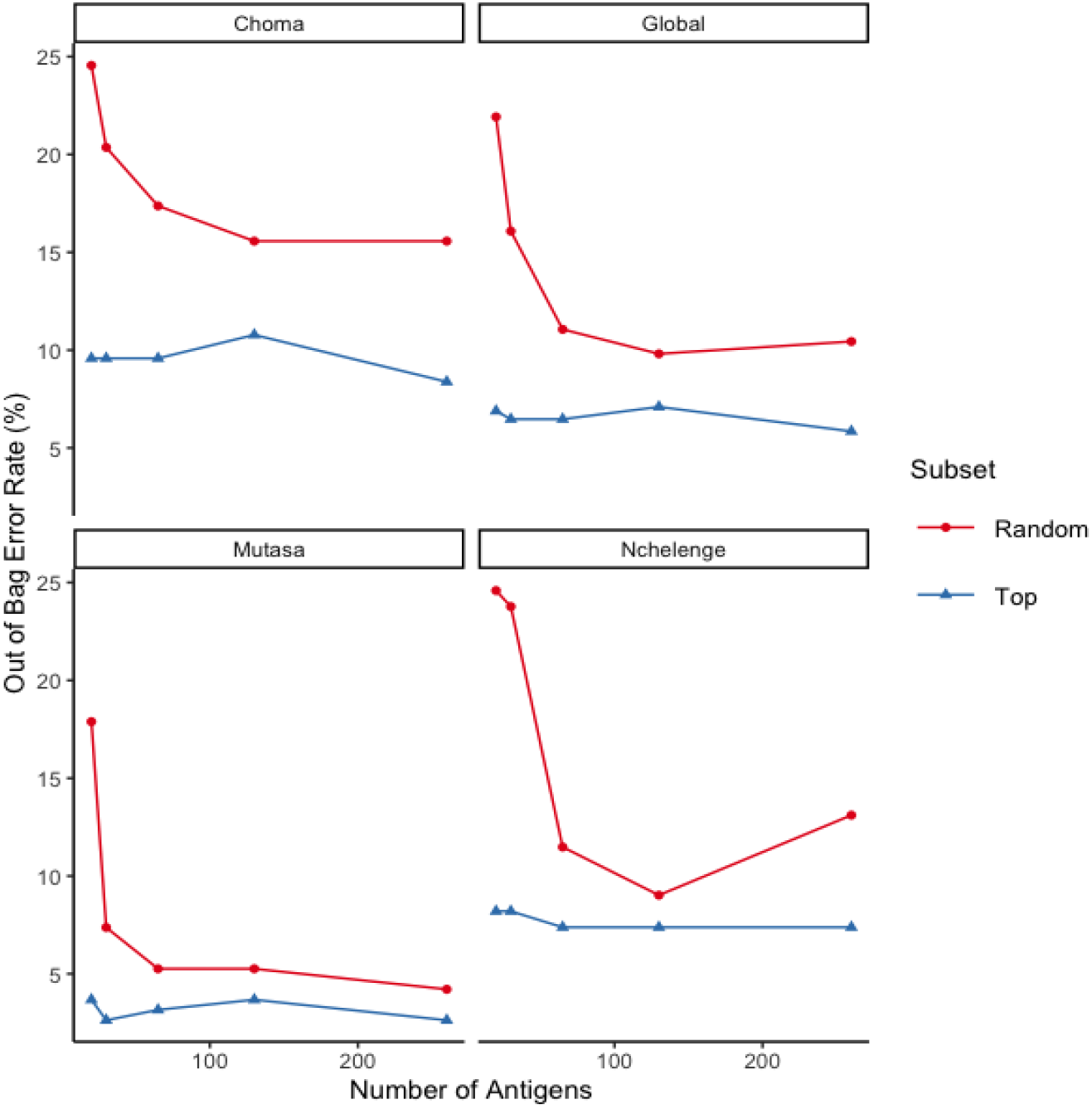
Out of bag error rate represented as a percentage for random forest models with 20, 30, 65, 130, and 260 antigens. Subsets of antigens are either the most important as determined by mean decrease in Gini index (Top, in blue), or a random subset (Random, in red). All models are fit to all 479 serum samples and out of bag error rates are reported for global, and category specific error rates (Choma, Nchlenege, or Mutasa).

### 4.2 Bioinformatic pipelines and analytic methods optimized for protein microarray data allow for an expanded analysis of biomarkers of past infection in adult populations

Elucidating markers of the timing and intensity of past exposure in adults is further complicated by uniformly high antibody responses to *P. falciparum* due to high numbers of re-exposure and re-infection events. However adults’ infection histories span longer periods and therefore contain important information about historical transmission trends that enable longitudinal surveillance. Random forest models fit to subsets of samples reveal that the ranks produced by the bioinformatic pipeline (Section 3) and analytic models that leverage interactions between predictors classify samples from adults (using an age cutoff of 5 years and of 15 years) into their region of origin with a global OOB error under 10% (Figure 2) and AUC values over 95% (Table S3). Although the class-specific OOB error rates are much more variable for this subset analysis (Figure 2), this is likely due to the relatively small sample sizes of some subsets, particularly children under age 5 years. Although a formal test-training split was not carried out for the subset analysis, the similar values of AUC in Tables S1 and S2 obtained for classification with all samples suggest that AUC values in Table S3 are robust. These results could indicate that differences in recent exposure lead to distinct antibody responses among adults and allow the random forest models to classify samples into regions of different historical transmission patterns with high accuracy. However, given the duration of adults’ exposure, it is also possible that longer lasting antibody responses are reflective of differences in historical exposure, and these are what enable accurate classification among samples from adults. This is further supported by the high accuracy of classification by these models among RDT negative individuals (global OOB error under 10% Figure 2 and AUC over 95% Table S3) who were unlikely to be harboring parasites in the weeks preceding sampling.

**Figure 2:**
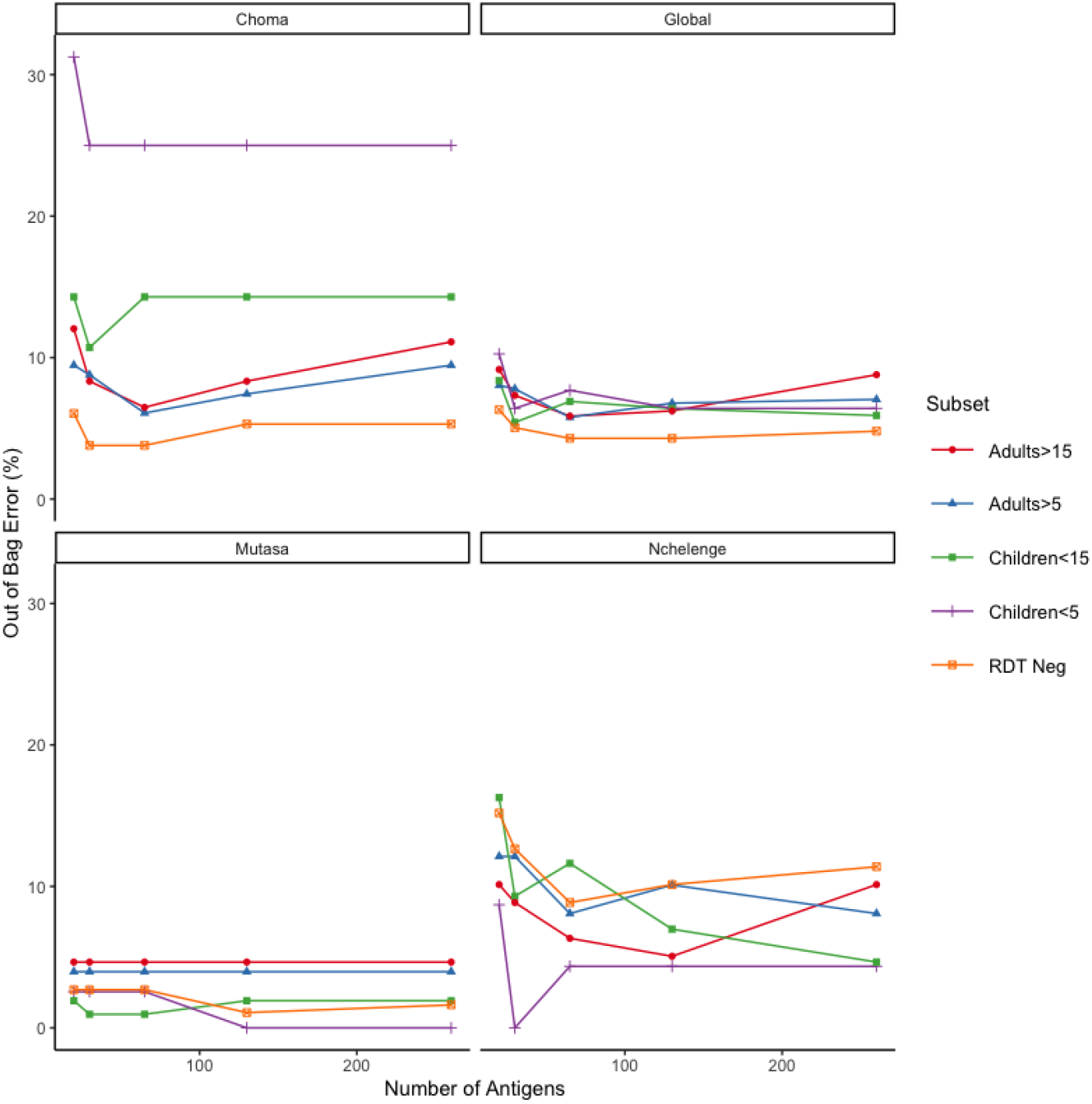
Out of bag error rate represented as a percentage for random forest models with the most important 20, 30, 65, 130, and 260 antigens as determined by mean decrease in Gini index. Random forest models are run on five subsets of samples: children under 5 (n=78), adults over 5 (n=398), children under 15 (n=203), adults over 15 (n=273),and RDT negative (n=396). Out of bag error rates are reported for global, and category specific error rates (Choma, Nchlenege, or Mutasa).

Using the antigen ranks from the bioinformatic pipeline (Section 3) as predictors, the random forest models outperformed polytomous, logistic regression models, the latter of which do not include interactions between the ranks of antibody responses to antigens. The difference between these models’ performance is most extreme for models that use the top (determined by mean decrease in Gini index in the random forest model) 130 and 260 antigens (Table 2).

**Table 2:**
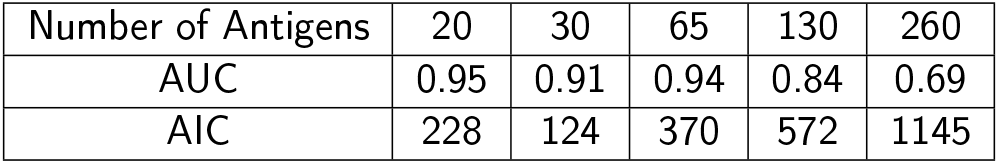
Area under the curve (AUC) and Akaike Information Criterion (AIC) for polytomous, logistic regression models. AUC is computed based on a 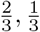 train-test split of samples. The AUC reported is a multi-class AUC computed with the macro-average method as described in Hand and Till (2001).

| Number of Antigens | 20 | 30 | 65 | 130 | 260 |
| --- | --- | --- | --- | --- | --- |
| AUC | 0.95 | 0.91 | 0.94 | 0.84 | 0.69 |
| AIC | 228 | 124 | 370 | 572 | 1145 |

These findings suggest that the interactions between ranked responses to antigens could help to distinguish the antibody responses to *Plasmodium* antigens of adults across zones of differing historical malaria transmission.

### 4.3 Most important antigens for classification reveal novel candidate biomarkers for the timing and intensity of past exposure to *P. falciparum*

Generally, there is little overlap between the biomarkers identified in previous studies aimed at identifying the intensity and timing of past exposure to *P. falciparum* and the most important antigens for classification via random forest identified here (as determined by mean decrease in Gini index). This suggests that analysis of these data has revealed novel candidate biomarkers for the intensity and timing of past exposure to *P. falciparum*. The 30 most important antigens (Table 3) contain a a large proportion 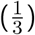 of *P. vivax* antigens despite no evidence that *P. vivax* circulated in any of the three study areas, although most of these proteins are designated as conserved suggesting they contain at least a motif of amino acids that is present across *Plasmodium* species, which may explain a certain level of cross-reactivity with antibodies generated after *P. falciparum* infection. Additionally, random forest models with only the top *P. falciparum* antigens (excluding *P. vivax* antigens as possible predictors) have similar OOB error to ones with the top *P. falciparum* and *vivax* antigens (Figure S2 and Table S5). Two of the top 30 antigens, Plasmodium exported protein (PHISTc), and serine repeat antigen 4(SERA4) overlap with published markers of exposure from Kobayashi *and others* (2019) and Helb *and others* (2015). Additionally, certain proteins have been characterized in other studies related to human antibody response to *P. falciparum* infection including: clustered-asparagine-rich protein which has been shown to be recognized by human T-cells and antibodies in recently exposed individuals (Wahlgren *and others*, 1991); merzoite surface protien 3 (MSP3) which has been shown to generate antibody response during natural *P. falciparum* infection (Deshmukh *and others*, 2018); erythrocyte binding antigen 175 (EBA175), which has been shown to elicit antibody responses in children that are protective against clinical malaria (Mccarra *and others*, 2011); and an erythrocyte membrane protein1, PfEMP1 (VAR), which appears as different proteins in the same family among the lists of biomarkers identified by Helb *and others* (2015) and Kobayashi *and others* (2019). Aside from these, several proteins in the list of 30 most important antigens for classification (Table 3) could be novel biomarkers of the intensity and timing past exposure to *P. falciparum*, and considering the list of 65 most important antigens (Table S4) reveals an expanded set of such candidate markers.

**Table 3.**
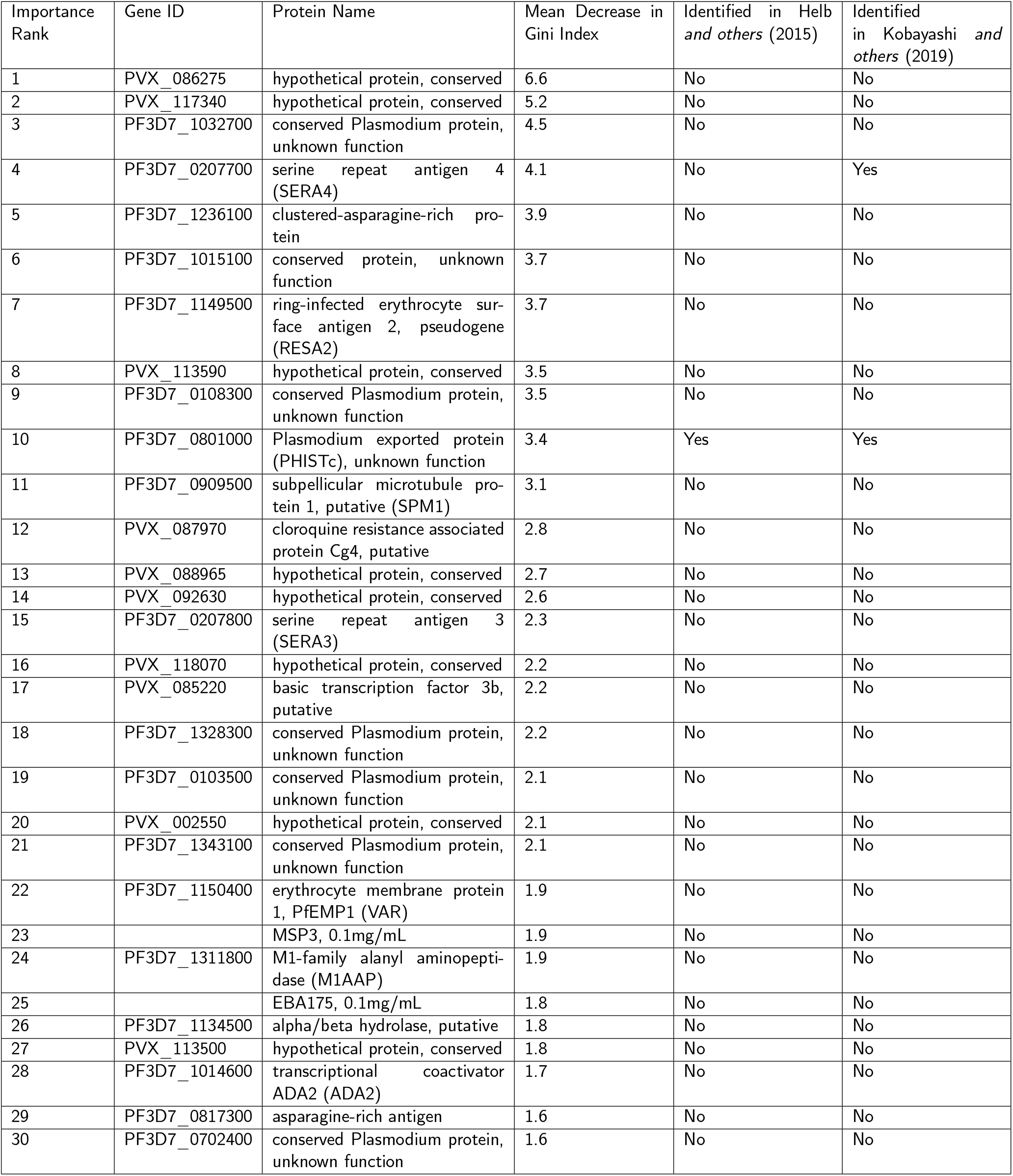
The 30 most important *Plasmodium falciparum* and *P. vivax* antigens for classification via random forest identified according to the mean decrease in Gini index. Overlaps with previously published results from Helb *and others* (2015) and Kobayashi *and others* (2019) are shown.

## 5 Discussion

Using protein microarrays, human antibody responses to *P. falciparum* and *P.vivax* antigens were characterized across serum samples collected from three regions in Zambia and Zimbabwe with historically high (Nchelenge District, Zamiba - parasite prevalence of 50%), intermediate (Mutasa District, Zimbabwe - parasite prevalence of 10%), and low levels (Choma District, Zambia - parasite prevalence of 1%) of malaria transmission (Moss *and others*, 2015; Mharakurwa *and others*, 2012).Previously, analysis performed using common pre-processing steps found an association between responses to the 30 most reactive antigens and the region of origin of a sample, but only among children under 5 years (Kobayashi *and others*, 2019). We expanded this analysis, and overcame some obstacles associated with finding markers of infection history in older adults by considering responses to more antigens and by ensuring a minimal amount of technical variation was left in the data. We accomplished the goals of minimizing between-sample technical variation and expanding the list of candidate antigens with the application of a novel bioinformatic pipeline (Bérubé *and others*, 2021, 2022). Briefly, we obtained ranks based on fluorescent intensity of each antigen within each array that represent an optimal estimate (Lin *and others*, 2006) of the relative level of antibody response of an individual to a particular antigen, as compared to the response of that same individual to other *Plasmodium* antigens. Using the ranks of between 20 and 260 of the antigens measured on the protein microarrays as predictors in a random decision forest we classified samples into their region of origin with high accuracy. A polytomous, logistic regression model did not classify samples with such high accuracy, suggesting that differences in individuals’ reactivity to these antigens, along with interactions among these predictors, is reflective of differences in past intensity and frequency of exposure to *P. falciparum*. Therefore, the antigens that classify samples into their region of origin are candidate biomarkers for the timing and intensity of past exposure to *P. falciparum*, and could overcome challenges associated with current surveillance methods.

Although the analysis we performed identified candidate biomarkers for the timing and intensity of past exposure, there are several additional studies and steps that need to be carried out before any biomarkers can be validated and used in the field for surveillance. A longitudinal study with detailed information about participant’s infection status over time, possibly spanning decades, would be required to directly associate differences in antibody response to these antigens with differences in infection history. Moreover, given the geographical heterogeneity in the genomes of *P. falciparum* parasites, it is possible that the antibody responses in the study participants in Zambia and Zimbabwe are not easily generalized to other populations; a more extensive geographical sample across sub-Saharan Africa is required. Furthermore, protein microarray data, even with careful pre-processing, remains prone to high levels of technical variation and does not allow analysts to directly measure antibody levels. Therefore, a lower throughput assay that does enable such measurements like a multiplex bead array or an enzyme-linked imunosorbent assay (ELISA) should be performed with the candidate antigens identified here to further validate their potential as biomarkers for the timing and intensity of past exposure to *P. falciparum*. Finally, the cross-validated and OOB error rates of models with sequentially fewer antigens as predictors suggest that 20 is sufficient to predict region of origin with high accuracy and there is little added predictive accuracy in models with up to 260 antigens, however, models with fewer than 10 antigens as predictors have substantially higher cross-validated and OOB error rates (Table S6).

The number of *P. vivax* antigens that are among the most important for classifying samples into their region of origin, 10 of the top 30 and 25 of the top 65, is surprising given that cases of *P. vivax* have not been and are not currently documented in any of the sampling regions. Many of the *P. vivax* antigens in list of top 30 (Table 3) or top 65 antigens (Table S4), are conserved proteins, meaning they contain at least a motif that is conserved across *Plasmodium* species. Therefore, this finding could be the result of antibodies generated in response to *P. falciparum* exposure that cross-react with *P. vivax* antigens. However, further investigation into the possible utility of including *P. vivax* antigens in the search for markers of the timing and intensity of past *P. falciparum* exposure may be worthwhile.

Although further studies are required to validate the candidate biomarkers we have identified in this study, narrowing down a set of possible antigens to target is an important first step toward eventually integrating serology into routine malaria surveillance. Ultimately, validated markers of the timing and intensity of past exposure to *P. falciparum* could enable analysts to accurately assess the burden of malaria over long periods of time with the ease of a cross-sectional sample. This, in turn could facilitate targeting effective interventions in low and high transmission settings and accurately evaluating the effectiveness of such interventions.

