## Supplement for "A Random Forest Classifier Uses Antibody Responses to *Plasmodium* Antigens to Reveal Candidate Biomarkers of the Intensity and Timing of Past Exposure to *Plasmodium falciparum*"

February 16, 2022

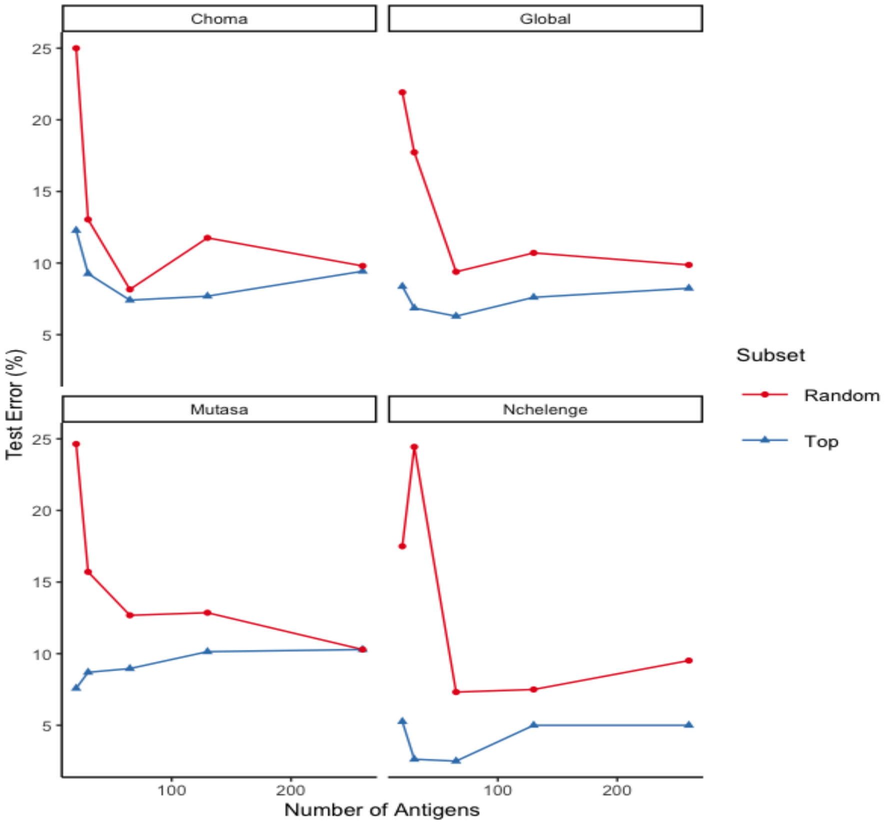


**Fig S 1:** Test error rate represented as a percentage for random forest models with 20, 30, 65, 130, and 260 antigens. Subsets of antigens are either the most important as determined by mean decrease in Gini index (Top, in blue), or a random subset (Random, in red). All models are fit to a random subset of 318 samples ($\frac{2}{3}$ of the serum samples) and test error rates (computed with the remaining $\frac{1}{3}$ of the serum samples) are reported for global, and category specific error rates (Choma, Nchlenege, or Mutasa).

**Table S 1:** Area under the curve and corresponding 95 % confidence intervals (CIs) for random forest models run with the top 20,30,65,130,and 260 antigens, as determined by mean decrease in Gini index. Models are fit with all 479 samples, and multi-class classification AUCs are computed using the the multiROC package in R[3] with the macro-average for the global AUC and a basic bootstrap to compute the 95% CIs.

| Number of Antigens | 20 | 30 | 65 | 130 | 260 |
|---|---|---|---|---|---|
| Choma | 0.98 (0.97,0.99) | 0.98 (0.97,0.99) | 0.99 (0.98,0.99) | 0.99 (0.98,0.99) | 0.98 (0.98,0.99) |
| Nchelenge | 0.99 (0.98,1.00) | 0.99 (0.99,1.00) | 0.99 (0.99,1.00) | 0.99 (0.99,1.00) | 0.99 (0.99,1.00) |
| Mutasa | 0.99 (0.99,1.00) | 0.99 (0.99,1.00) | 1.00 (0.99,1.00) | 0.99 (0.99,1.00) | 1.00 (0.99,1.00) |
| Global | 0.99 (0.98,0.99) | 0.99 (0.98,1.00) | 0.99 (0.99,1.00) | 0.99 (0.99,1.00) | 0.99 (0.99,1.00) |

**Table S 2:** Area under the curve and corresponding 95 % confidence intervals (CIs) for random forest models run with the top 20,30,65,130,and 260 antigens, as determined by mean Gini index. Models are fit with a random subset of 318 samples ($\frac{2}{3}$ of the serum samples) and AUC values are computed for the remaining 161 ($\frac{1}{3}$) serum samples. Global, and category specific error rates (Choma, Nchlenege, or Mutasa) are reported using multi-class classification AUCs calculated using the the multiROC package in R[3] with the macro-average for the global AUC and a basic bootstrap to compute the 95% CIs.

| Number of Antigens | 20 | 30 | 65 | 130 | 260 |
|---|---|---|---|---|---|
| Choma | 0.98(0.96,1.00) | 0.98 (0.96,1.00) | 0.98 (0.97,1.00) | 0.98 (0.97,1.00) | 0.98 (0.97,1.00) |
| Nchelenge | 0.98 (0.97,1.00) | 0.98 (0.97,1.00) | 0.99 (0.97,1.00) | 0.99 (0.99,1.00) | 0.99 (0.99,1.00) |
| Mutasa | 1.00 (0.99,1.00) | 0.99 (0.99,1.00) | 1.00 (0.99,1.00) | 1.00 (0.99,1.00) | 1.00 (0.99,1.00) |
| Global | 0.98 (0.97,1.00) | 0.98 (0.97,1.00) | 0.99 (0.98,1.00) | 0.99 (0.98,1.00) | 0.99 (0.98,1.00) |

**Table S 3:** Area under the curve and corresponding 95 % confidence intervals (CIs) for random forest models run with the top 20,30,65,130,and 260 antigens, as determined by mean derease in Gini index. Models are fit with five subsets of samples: children under 5 (n=78), adults over 5 (n=398), children under 15 (n=203), adults over 15 (n=273),and RDT negative (n=396). Multi-class classification AUCs are computed using the the multiROC package in R[3] with the macro-average for the global AUC and a basic bootstrap to compute the 95% CIs.

| Group | Error Type | 20 | 30 | 65 | 130 | 260 |
|---|---|---|---|---|---|---|
| Children $\leq 5$ | Global | 0.98 (0.97, 1.00) | 0.99 (0.98, 1.00) | 0.99 (0.98, 1.00) | 0.99 (0.98, 1.00) | 0.98 (0.96, 1.00) |
| | Choma | 0.97 (0.95, 1.00) | 0.99 (0.97, 1.00) | 0.98 (0.96, 1.00) | 0.98 (0.96, 1.00) | 0.95 (0.91, 1.00) |
| | Nchelenge | 0.99 (0.98, 1.00) | 1.00 (0.99, 1.00) | 1.00 (1.00, 1.00) | 1.00 (1.00, 1.00) | 1.00 (1.00, 1.00) |
| | Mutasa | 1.00 (0.99, 1.00) | 1.00 (0.99, 1.00) | 1.00 (1.00, 1.00) | 1.00 (1.00, 1.00) | 1.00 (0.99, 1.00) |
| Adults $> 5$ | Global | 0.98 (0.97, 0.99) | 0.98 (0.98, 0.99) | 0.99 (0.98, 1.00) | 0.99 (0.98, 1.00) | 0.99 (0.98, 0.99) |
| | Choma | 0.97 (0.96, 0.99) | 0.97 (0.96, 0.99) | 0.98 (0.97, 0.99) | 0.98 (0.97, 0.99) | 0.98 (0.97, 0.99) |
| | Nchelenge | 0.99 (0.98, 1.00) | 0.99 (0.98, 1.00) | 0.99 (0.98, 1.00) | 0.99 (0.99, 1.00) | 0.99 (0.99, 1.00) |
| | Mutasa | 0.99 (0.98, 1.00) | 0.99 (0.98, 1.00) | 0.99 (0.99, 1.00) | 0.99 (0.98, 1.00) | 0.99 (0.98, 1.00) |
| Children $\leq 15$ | Global | 0.99 (0.98, 1.00) | 0.99 (0.98, 1.00) | 0.99 (0.98, 1.00) | 0.99 (0.98, 1.00) | 0.99 (0.98, 1.00) |
| | Choma | 0.98 (0.97, 1.00) | 0.98 (0.97, 1.00) | 0.98 (0.97, 1.00) | 0.98 (0.96, 1.00) | 0.98 (0.97, 1.00) |
| | Nchelenge | 0.99 (0.98, 1.00) | 0.99 (0.98, 1.00) | 0.99 (0.98, 1.00) | 0.99 (0.99, 1.00) | 0.99 (0.99, 1.00) |
| | Mutasa | 0.99 (0.99, 1.00) | 1.00 (0.99, 1.00) | 0.99 (0.99,1.00) | 0.99 (0.99, 1.00) | 0.99 (0.99, 1.00) |
| Adults $> 15$ | Global | 0.98 (0.96, 0.99) | 0.98 (0.97, 1.00) | 0.99 (0.98, 1.00) | 0.99 (0.98, 1.00) | 0.98 (0.97, 1.00) |
| | Choma | 0.96 (0.94, 0.98) | 0.97 (0.96, 0.99) | 0.98 (0.97, 1.00) | 0.98 (0.97, 1.00) | 0.97 (0.96, 0.99) |
| | Nchelenge | 0.98 (0.97, 1.00) | 0.99 (0.98, 1.00) | 0.99 (0.99, 1.00) | 1.00 (0.99, 1.00) | 0.99 (0.99, 1.00) |
| | Mutasa | 0.99 (0.97, 1.00) | 0.99 (0.98, 1.00) | 0.99 (0.97, 1.00) | 0.98 (0.97, 1.00) | 0.98 (0.96, 1.00) |
| RDT Neg | Global | 0.99 (0.98, 1.00) | 0.99 (0.98, 1.00) | 0.99 (0.99, 1.00) | 0.99 (0.99, 1.00) | 0.99 (0.99, 1.00) |
| | Choma | 0.98 (0.97, 0.99) | 0.99 (0.98, 1.00) | 0.99 (0.98, 1.00) | 0.99 (0.98, 1.00) | 0.99 (0.98, 1.00) |
| | Nchelenge | 0.98 (0.98, 1.00) | 0.99 (0.98, 1.00) | 0.99 (0.98, 1.00) | 0.99 (0.98, 1.00) | 0.99 (0.98, 1.00) |
| | Mutasa | 1.00 (0.98, 1.00) | 0.99 (0.98, 1.00) | 1.00 (0.99, 1.00) | 1.00 (0.99, 1.00) | 1.00 (0.99, 1.00) |

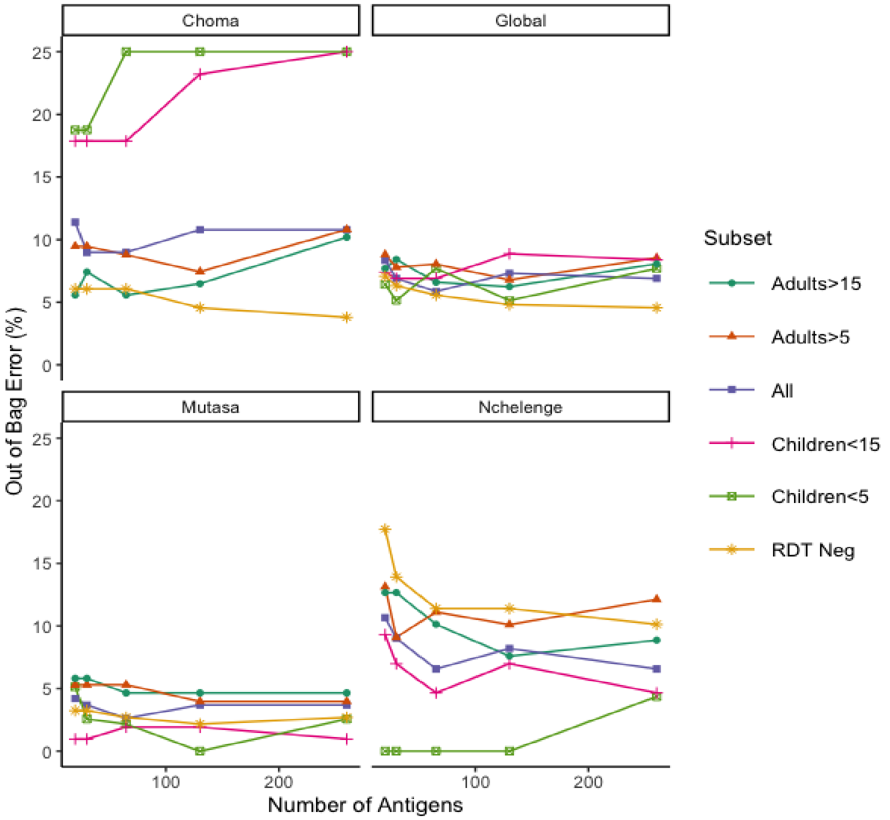


**Fig S 2:** Out of bag error rate represented as a percentage for random forest models with the most important 20, 30, 65, 130, and 260 *P. falciparum* antigens. Antigen importance is determined by mean decrease in Gini index. All models are fit to five subsets of serum samples: children under 5 (n=78), adults over 5 (n=398), children under 15 (n=203), adults over 15 (n=273),and individuals who are RDT negative at the time of sampling (n=396), as well as all 479 samples. Errors are reported for global, and category specific error rates (Choma, Nchlenege, or Mutasa).

**Table S 4:** The 65 most important *Plasmodium falciparum* and *P. vivax* antigens for classification via random forest identified according to the mean decrease in Gini index. Overlaps with previously published results from Helb *and others*[1] and Kobayashi *and others*[2] are shown.

| Importance Rank | Gene ID | Protein Name | Mean Decrease in Gini Index | Identified in Helb *and others*[1] | Identified in Kobayashi *and others*[2] |
|---|---|---|---|---|---|
| 1 | PVX_086275 | hypothetical protein, conserved | 6.6 | No | No |
| 2 | PVX_117340 | hypothetical protein, conserved | 5.2 | No | No |
| 3 | PF3D7_1032700 | conserved Plasmodium protein, unknown function | 4.5 | No | No |
| 4 | PF3D7_0207700 | serine repeat antigen 4 (SERA4) | 4.1 | No | Yes |
| 5 | PF3D7_1236100 | clustered-asparagine-rich protein | 3.9 | No | No |
| 6 | PF3D7_1015100 | conserved protein, unknown function | 3.7 | No | No |
| 7 | PF3D7_1149500 | ring-infected erythrocyte surface antigen 2, pseudogene (RESA2) | 3.7 | No | No |
| 8 | PVX_113590 | hypothetical protein, conserved | 3.5 | No | No |
| 9 | PF3D7_0108300 | conserved Plasmodium protein, unknown function | 3.5 | No | No |
| 10 | PF3D7_0801000 | Plasmodium exported protein (PHISTc), unknown function | 3.4 | Yes | Yes |
| 11 | PF3D7_0909500 | subpellicular microtubule protein 1, putative (SPM1) | 3.1 | No | No |
| 12 | PVX_087970 | cloroquine resistance associated protein Cg4, putative | 2.8 | No | No |
| 13 | PVX_088965 | hypothetical protein, conserved | 2.7 | No | No |
| 14 | PVX_092630 | hypothetical protein, conserved | 2.6 | No | No |
| 15 | PF3D7_0207800 | serine repeat antigen 3 (SERA3) | 2.3 | No | No |
| 16 | PVX_118070 | hypothetical protein, conserved | 2.2 | No | No |
| 17 | PVX_085220 | basic transcription factor 3b, putative | 2.2 | No | No |
| 18 | PF3D7_1328300 | conserved Plasmodium protein, unknown function | 2.2 | No | No |

Continued on next page

**Table 4 –continued from previous page**

| Importance Rank | Gene ID | Protein Name | Mean Decrease in Gini Index | Identified in Helb *and others*[1] | Identified in Kobayashi *and others*[2] |
|---|---|---|---|---|---|
| 19 | PF3D7_0103500 | conserved Plasmodium protein, unknown function | 2.1 | No | No |
| 20 | PVX_002550 | hypothetical protein, conserved | 2.1 | No | No |
| 21 | PF3D7_1343100 | conserved Plasmodium protein, unknown function | 2.1 | No | No |
| 22 | PF3D7_1150400 | erythrocyte membrane protein 1, PfEMP1 (VAR) | 1.9 | No | No |
| 23 | NA | MSP3, 0.1mg/mL | 1.9 | No | No |
| 24 | PF3D7_1311800 | M1-family alanyl aminopeptidase (M1AAP) | 1.9 | No | No |
| 25 | NA | EBA175, 0.1mg/mL | 1.8 | No | No |
| 26 | PF3D7_1134500 | alpha/beta hydrolase, putative | 1.8 | No | No |
| 27 | PVX_113500 | hypothetical protein, conserved | 1.8 | No | No |
| 28 | PF3D7_1014600 | transcriptional coactivator ADA2 (ADA2) | 1.7 | No | No |
| 29 | PF3D7_0817300 | asparagine-rich antigen | 1.6 | No | No |
| 30 | PF3D7_0702400 | conserved Plasmodium protein, unknown function | 1.6 | No | No |
| 31 | PVX_100910 | hypothetical protein, conserved | 1.6 | No | No |
| 32 | PVX_089790 | hypothetical protein | 1.5 | No | No |
| 33 | PVX_083040 | hypothetical protein, conserved | 1.4 | No | No |
| 34 | PF3D7_1036900 | conserved Plasmodium protein, unknown function | 1.4 | No | No |
| 35 | PF3D7_1477600 | surface-associated interspersed protein 14.1 (SURFIN 14.1) (Surf14.1) | 1.4 | No | No |
| 36 | PVX_091785 | translation elongation factor EF-1, subunit alpha, putative | 1.3 | No | No |
| 37 | PF3D7_1335100 | merozoite surface protein 7 (MSP7) | 1.3 | No | No |
| 38 | PF3D7_0501100 | heat shock protein 40, type II (HSP40) | 1.3 | Yes | No |
| 39 | PF3D7_1001600 | alpha/beta hydrolase, putative | 1.2 | No | No |

Continued on next page

**Table 4 –continued from previous page**

| Importance Rank | Gene ID | Protein Name | Mean Decrease in Gini Index | Identified in Helb *and others*[1] | Identified in Kobayashi *and others*[2] |
|---|---|---|---|---|---|
| 40 | PVX_097810 | hypothetical protein, conserved | 1.2 | No | No |
| 41 | PVX_087865 | hypothetical protein, conserved | 1.2 | No | No |
| 42 | PVX_100845 | hypothetical protein, conserved | 1.2 | No | No |
| 43 | NA | histone acetyltransferase GCN5 (GCN5) | 1.2 | No | No |
| 44 | PF3D7_1030800 | calmodulin, putative | 1.2 | No | No |
| 45 | PF3D7_1109200 | conserved Plasmodium protein, unknown function | 1.1 | No | No |
| 46 | PF3D7_1227100 | DNA helicase 60 (DH60) | 1.1 | No | No |
| 47 | PF3D7_0424800 | Plasmodium exported protein (PHISTb), unknown function | 1.1 | No | No |
| 48 | PF3D7_0713900 | conserved Plasmodium protein, unknown function | 1.1 | No | No |
| 49 | PVX_114445 | pyruvate kinase, putative | 1.1 | No | No |
| 50 | PF3D7_1433500 | DNA topoisomerase II, putative | 1.1 | No | No |
| 51 | PVX_091845 | ethanolamine kinase, putative | 1.1 | No | No |
| 52 | PVX_086010 | hypothetical protein, conserved | 1.0 | No | No |
| 53 | PF3D7_1020300 | cytoplasmic dynein intermediate chain, putative | 1.0 | No | No |
| 54 | PVX_095185 | hypothetical protein, conserved | 1.0 | No | No |
| 55 | PF3D7_0511600 | conserved Plasmodium protein, unknown function | 1.0 | No | No |
| 56 | PF3D7_0904900 | Cu2 -transporting ATPase, putative (CUP) | 1.0 | No | Yes |
| 57 | PVX_000820 | flap exonuclease, putative | 1.0 | No | No |
| 58 | PVX_090090 | CW-type zinc finger domain-containing protein | 1.0 | No | No |
| 59 | PF3D7_0713100 | Pfmc-2TM Maurer's cleft two transmembrane protein (MC-2TM) | 1.0 | No | No |

Continued on next page

**Table 4 –continued from previous page**

| Importance Rank | Gene ID | Protein Name | Mean Decrease in Gini Index | Identified in Helb *and others*[1] | Identified in Kobayashi *and others*[2] |
|---|---|---|---|---|---|
| 60 | PVX_085570 | hypothetical protein, conserved | 0.9 | No | No |
| 61 | PF3D7_1441800 | SNF7 family protein, putative | 0.9 | No | No |
| 62 | PF3D7_1007700 | transcription factor with AP2 domain(s) (ApiAP2) | 0.9 | No | Yes |
| 63 | PVX_004537 | hypothetical protein, conserved | 0.9 | No | No |
| 64 | PF3D7_1106300 | exonuclease, putative | 0.9 | Yes | No |
| 65 | PF3D7_1448000 | U3 snoRNA-associated small subunit rRNA processing protein, putative | 0.9 | No | No |

**Table S 5:** The 30 most important *Plasmodium falciparum* antigens for classification via random forest identified according to the mean decrease in Gini index. Overlaps with previously published results from Helb *and others*[1] and Kobayashi *and others*[2] are shown.

| Importance Rank | Gene ID | Protein Name | Mean Decrease in Gini Index | Identified in Helb *and others*[1] | Identified in Kobayashi *and others*[2] |
|---|---|---|---|---|---|
| 1 | PF3D7_1032700 | conserved Plasmodium protein, unknown function | 6.7 | No | No |
| 2 | PF3D7_1015100 | conserved protein, unknown function | 6.6 | No | No |
| 3 | PF3D7_1236100 | clustered-asparagine-rich protein | 5.6 | No | No |
| 4 | PF3D7_0207700 | serine repeat antigen 4 (SERA4) | 5.8 | No | Yes |
| 5 | PF3D7_1149500 | ring-infected erythrocyte surface antigen 2, pseudogene (RESA2) | 4.9 | No | No |
| 6 | PF3D7_0801000 | Plasmodium exported protein (PHISTc), unknown function | 4.2 | Yes | Yes |
| 7 | PF3D7_0108300 | conserved Plasmodium protein, unknown function | 4.1 | No | No |
| 8 | PF3D7_0909500 | subpellicular microtubule protein 1, putative (SPM1) | 4.0 | No | No |
| 9 | PF3D7_1134500 | alpha/beta hydrolase, putative | 3.7 | No | No |
| 10 | NA | MSP3, 0.1mg/mL | 3.4 | No | No |
| 11 | PF3D7_1311800 | M1-family alanyl aminopeptidase (M1AAP) | 3.3 | No | No |
| 12 | PF3D7_1343100 | conserved Plasmodium protein, unknown function | 3.2 | No | No |
| 13 | PF3D7_1106300 | exonuclease, putative | 2.9 | Yes | No |
| 14 | PF3D7_0501100 | heat shock protein 40, type II (HSP40) | 2.8 | Yes | No |
| 15 | PF3D7_0207800 | serine repeat antigen 3 (SERA3) | 2.8 | No | No |
| 16 | PF3D7_0103500 | conserved Plasmodium protein, unknown function | 2.8 | No | No |
| 17 | NA | EBA175, 0.1mg/mL | 2.7 | No | No |
| 18 | PF3D7_1014600 | transcriptional coactivator ADA2 (ADA2) | 2.7 | No | No |
| 19 | PF3D7_1036900 | conserved Plasmodium protein, unknown function | 2.6 | No | No |
| 20 | PF3D7_1477600 | surface-associated interspersed protein 14.1 (SURFIN 14.1) (Surf14.1) | 2.5 | No | No |
| 21 | PF3D7_1328300 | conserved Plasmodium protein, unknown function | 2.4 | No | No |
| 22 | PF3D7_1007700 | transcription factor with AP2 domain(s) (ApiAP2) | 2.3 | No | Yes |
| 23 | NA | histone acetyltransferase GCN5 (GCN5) | 2.3 | No | No |
| 24 | PF3D7_0511600 | conserved Plasmodium protein, unknown function | 2.3 | No | No |
| 25 | PF3D7_0702400 | conserved Plasmodium protein, unknown function | 2.2 | No | No |
| 26 | PF3D7_1479000 | acyl-CoA synthetase (ACS1a) | 2.2 | No | No |
| 27 | PF3D7_1109200 | conserved Plasmodium protein, unknown function | 2.2 | No | No |
| 28 | PF3D7_1335100 | merozoite surface protein 7 (MSP7) | 2.1 | No | No |
| 29 | PF3D7_0424800 | Plasmodium exported protein (PHISTb), unknown function | 2.0 | No | No |
| 30 | NA | Pf LSA1, 0.3mg/mL | 2.0 | No | No |

**Table S 6:** Cross-validated and OOB error rates for random forest models with sequentially fewer antigens as predictors. In each model with sequentially fewer predictors, the antigens retained in the model are those with the highest values of mean decrease in Gini index as originally computed in the random forest model with all 1038 antigens as predictors.

| Number of Antigens | 1038 | 519 | 260 | 130 | 65 | 32 | 16 | 8 | 4 | 2 |
|---|---|---|---|---|---|---|---|---|---|---|
| Cross-validated Error (%) | 7.5 | 6.9 | 6.5 | 6.3 | 6.1 | 7.1 | 7.9 | 13.1 | 23.0 | 37.4 |
| OOB Error (%) | 7.7 | 7.5 | 6.3 | 5.6 | 5.9 | 6.5 | 7.9 | 15.9 | 23.6 | 35.9 |
